# Genome sequence of tsetse bracoviruses: insights into symbiotic virus evolution

**DOI:** 10.1101/045088

**Authors:** Kelvin M. Kimenyi, Muna F. Abry, Winnie Okeyo, Enock Matovu, Daniel Masiga, Benard W. Kulohoma

**Affiliations:** Center for Biotechnology and Bioinformatics, University of Nairobi P.O. Box 30197, 00100, Nairobi, Kenya; International Centre of Insect Physiology and Ecology, P.O. Box 30772, 00100, Nairobi, Kenya; Makerere University, P.O. Box 7062, Kampala, Uganda

**Keywords:** bracoviruses, endogenous viruses, eukaryotes

## Abstract

Mutualism between endogenous viruses and eukaryotes is still poorly understood. Whole genome data has highlighted the diverse distribution of viral sequences in several eukaryote host genomes. A group of endogenous double-stranded polydnaviruses known as bracoviruses has been identified in parasitic braconid wasp (Hymenoptera). Bracoviruses allow wasps to reproductively co-opt other insect larvae. Bracoviruses are excised from the host genome and injected in to the larva along side the wasp eggs; where they encode proteins that lower host immunity allowing development of parasitoid wasp larvae in the host. Interestingly, putative bracoviral sequences have recently been detected in the first sequenced genome of the tsetse fly (Diptera). This is peculiar since tsetse flies do not share this reproductive lifestyle. To investigate genome rearrangements associated with these unique mutual symbiotic relationships and examine its value as a potential vector control strategy entry point. We use comparative genomics to determine the presence, prevalence and genetic diversity of bracoviruses of five tsetse fly species (*G. austeni, G. brevipalpis, G. f. fuscipes, G. m. morsitans* and *G. pallidipes*) and the housefly (*Musca domestica*). We identify and use four viral Maverick genes as evolutionary models for bracoviruses. This is the first record of homologous bracoviruses in multiple *Dipteran* genomes. Phylogenetic reconstruction of each gene revealed two major clades that represent the two types of Mavericks. We detect varying magnitudes of purifying selection across these loci except for the poxvirus A32 gene, which is under positive selection. Moreover, these genes were inserted at conserved regions and co-evolve at similar rates with the host genomes.

## Background

Mutualism between eukaryotes and viruses is rare, since most viruses have parasitic associations with their hosts (Federici and Bigot 2003; Espagne et al. 2004; Burke et al. 2014). A group of double-stranded DNA (dsDNA) viruses called polydnaviruses (PDVs) have symbiotic associations with thousands of parasitoid wasps (order *Hymenoptera*), which parasitize other immunocompetent lepidopteran larvae to enable successful reproduction (Strand and Burke 2012). PDVs have co-evolved with wasps and present a unique opportunity to investigate genome rearrangements associated with these unique mutual symbiotic relationships (Espagne et al. 2004; Jancek et al. 2013). PDVs are broadly classified into two distinctly evolved genera: *Bracovirus* and *Ichnovirus* (Gundersen-Rindal and Pedroni 2006; Bezier et al. 2013). Bracoviruses are common within a monophyletic group of wasps known as the microgastroid complex, and are descendants of nudiviruses, a sister group of baculoviruses (Theze et al. 2011). It is estimated that bracoviruses were first integrated into the genome of the ancestor wasp approximately 100 million years ago (mya) (Herniou et al. 2013). Mutualism between wasps and bracoviruses developed over time, and involved genes of both wasp and viral origin. This functional association is estimated to date back to around 73.7±10 mya (Whitfield 2002).

Bracoviruses exist in two forms: a linear provirus integrated into the host genome that mediates vertical transmission as Mendelian traits, and virions containing circular dsDNA (Desjardins et al. 2007; Bezier et al. 2009). Viral replication, particle production and packaging into virions occur exclusively in a specialized part of the wasp ovaries (the calyx) (Bezier et al. 2009), and precede injection alongside one or more wasp eggs into the parasitized caterpillar host during wasp oviposition (Louis et al. 2013). Virions are replication deficient and their dsDNA is only expressed by the caterpillar host’s cellular replication machinery (Bezier et al. 2013; Chevignon et al. 2014). The virion particles encode proteins that compromise the caterpillar host immune defenses thus preventing recognition, encapsulation and destruction of the parasitoid eggs and larvae (Drezen et al. 2006; Herniou et al. 2013). However, lack of genes that independently encode viral structural proteins has elicited a debate on whether bracoviruses are of viral origin or are a ‘genetic secretion’ of the wasps (Bezier et al. 2009). An example is the bracoviral virion DNA in the wasp *Cotesia congregata* that consists of cellular genes of wasp origin, several viral genes and transposable elements (Drezen et al. 2006). Phylogenetic analysis of its functional bracoviral genes has highlighted sugar transporters of wasp origin (Desjardins et al. 2007). Transfer of these wasp genes into the provirus was facilitated by transposable elements, and subsequently followed by co-evolution with the host’s genome, to become more specialized (Espagne et al. 2004; Herniou et al. 2013; Jancek et al. 2013).

The recently released whole-genome sequence of the tsetse fly, *Glossina morsitans morsitans* (order *Diptera*), has revealed numerous putative bracoviral genes, which lack dipteran homologs, widely spread across the entire genome; in addition to a large DNA hytrosavirus, the *Glossina pallidipes* salivary gland hypertrophy virus (GpSGHV) (International Glossina Genome Initiative 2014). Although GpSGHV has been associated with reduced fecundity, life span, and causes salivary gland pathology in *Glossina*, its value as a potential entry point as a tsetse fly control strategy is untested. Perhaps more interesting is the finding of bracoviral sequences that bear close similarity (Basic Local Alignment Search Tool (BLAST), E values of <1e-50) to those identified in the parasitic wasps (order Hymenoptera) Glyptapanteles flavicoxis and Cotesia congregata, where they occur as PDVs (Espagne et al. 2004; International Glossina Genome Initiative 2014). Although the role of PDVs is well characterized in parasitic wasps, their organization, composition and functions in the tsetse fly genome is not known; indeed their presence is new information. Molecular dating estimates that the orders Diptera (includes the tsetse fly and the house fly) and Hymenoptera (includes wasps) diverged ~350 mya (Theze et al. 2011), which is prior to the estimated date of first integration of bracoviruses into the ancestor wasp genome (Herniou et al. 2013). This raises the possibility that these genes may be remnants of PDVs acquired before this separation, and tsetse flies lost bracoviral mutualism after they recently adapted to larviparity (development of a single larva in its uterus as opposed to laying multiple eggs). An alternate hypothesis is that an undetermined braconid wasp may have parasitized the tsetse fly ancestor (International Glossina Genome Initiative 2014).

Tsetse flies are important vectors that transmit African trypanosomiasis to humans (sleeping sickness) and cattle (nagana) (International Glossina Genome Initiative 2014). Approximately 70 million people and 50 million cattle are at risk of disease in tsetse-fly infested areas (Benoit et al. 2015). There are limited strategies of trypanosomiasis management primarily resulting from undesirable side effects of trypanocidal drug treatments; and there are emerging reports of multi-drug resistance (Barrett et al. 2007; Brun et al. 2010). According to the Pan African Tsetse and Trypanosomiasis Eradication Campaign (PATTEC), eradicating tsetse populations is the most viable approach of controlling trypanosomiasis in sub-Sahara Africa (Solano et al. 2010). Identification of *Glossina* genes regulating vectorial capacity is thus a priority, as their manipulation would provide important clues for the development of effective vector control strategies, which will greatly facilitate trypanosomiasis control (Barrett et al. 2007).

The tsetse fly, unlike other members of the order Diptera, does not lay eggs, but bears a fully developed larva (obligate adenotrophic viviparity) (Benoit et al. 2015). This makes it challenging to study PDVs in *Glossina* since during tsetse fly reproduction they are not replicated, excised from the host insect genome and packaged into viral particles that are mixed with semen like in parasitoid wasps. Wasp PDVs can easily be studied by first specifically extracting viral particles from the host (Bezier et al. 2013). Moreover, most tsetse fly bracoviruses consist of genes of host cellular origin with protein domains conserved across metazoans further complicating analysis (Bezier et al. 2008). We confirmed this by homology comparisons with the more recently sequenced dipteran genome of the house fly genome (Musca domestica), which is more closely related to the tsetse fly. Current homology searches of the same genes in the publicly accessible gene repository GenBank reveal homologues across Diptera (Supplementary Table 1).

Successful delivery of bracoviral genes by wasps into lepidopteran larvae has inspired notable agricultural applications. Currently, TSP14-producing transgenic plants effectively reduce *Manduca sexta* growth and development, thus protecting the plants from insect damage (Maiti et al. 2003). We investigate the prevalence, population genetics of bracoviruses across 5 *Glossina* species (*Glossina austeni, G. brevipalpis, G. fuscipes fuscipes* and *G. pallidipes*) genomes, assess whether the viral DNA is genetically active and identify clues to its function, compare genetic diversity relative to wasp bracoviruses, and try to deduce its origin. This newfound knowledge provides better understanding of tsetse biology, and novel possible intervention strategies. We show that that bracoviruses are present across the order Diptera and share close homology. In addition we identify the co-occurrence of two major bracoviral groups that co-evolve with the *Dipteran* host genome.

## Results

### Orthology between housefly and tsetse fly bracoviral sequences

We aimed to identify and compare sequence diversity of polydnaviruses (PDVs) in the recently sequenced genomes of five tsetse flies (*Glossina* species) and the housefly (*Musca domestica*). Previous work identified putative bracoviral homologues (n=310) in *G. m. morsitans* (International Glossina Genome Initiative 2014). In our analysis, we show that most of these bracoviral sequences have orthologs in other *Glossina* species and *M. domestica* (tBLASTn E-value <1e-50). In this dataset, most double stranded endogenous viral sequences identified have domains that are similar to those ubiquitously present in the host insect genome, and other metazoans in general. An example is the protein tyrosine phosphatase gene family, which plays a key role in regulating signal transduction pathways, and viral proteins thereby altering host physiology during parasitism (Espagne et al. 2004). This complicates unbiased selection and detailed analysis of bracoviral evolutionary biology. We circumvent this challenge by focusing on PDV sequences with virus-related domains that are good evolutionary biology models (Pritham et al. 2007) and are distinctly viral genes that act as distinguishing characters of the virus state (Koonin et al. 2006). In this work, we identify and use transposable elements designated Mavericks (Herniou et al. 2013) to underpin the genetic diversity of PDVs in the recently sequenced Dipteran genomes. Our Maverick sequences consist of four genes: a DNA polymerase B2 involved in DNA excision repair and initiation of replication (DNA Pol B 2) (Drezen et al. 2006), the N terminal region of the parvovirus coat protein VP1 (Parvo coat N) that is important for virion retention and transduction by insect vectors (Spitzer et al. 1997); a retroviral-like integrase; and poxvirus A32 protein (Pox A32) that encodes an ATPase involved in virion DNA packaging (Cassetti et al. 1998). Maverick associated genes have been shown to occur in close proximity to each other, and are thought to be genetically linked (Dupuy et al. 2011). We detected multiple Maverick associated genes in the different *Dipteran* genomes (Figure 1A). *G. austeni* has the highest number of Maverick associated genes (n=18) and *G. brevipalpis* the least (n=6). We also observed that parvo coat N was the most abundant gene across species (n=22), while the retroviral-like integrase was the least abundant (n=12), across all species. These bracoviral genes are widely distributed in the respective species genomes (Supplementary Table 1), and vary in length (Figure 1B). The parvo coat N gene has the most conserved length across all species, and there is remarkable variation at the other loci, emphasizing the importance of understanding the role of selection on genetic diversity at bracoviral loci.

**Figure 1.**
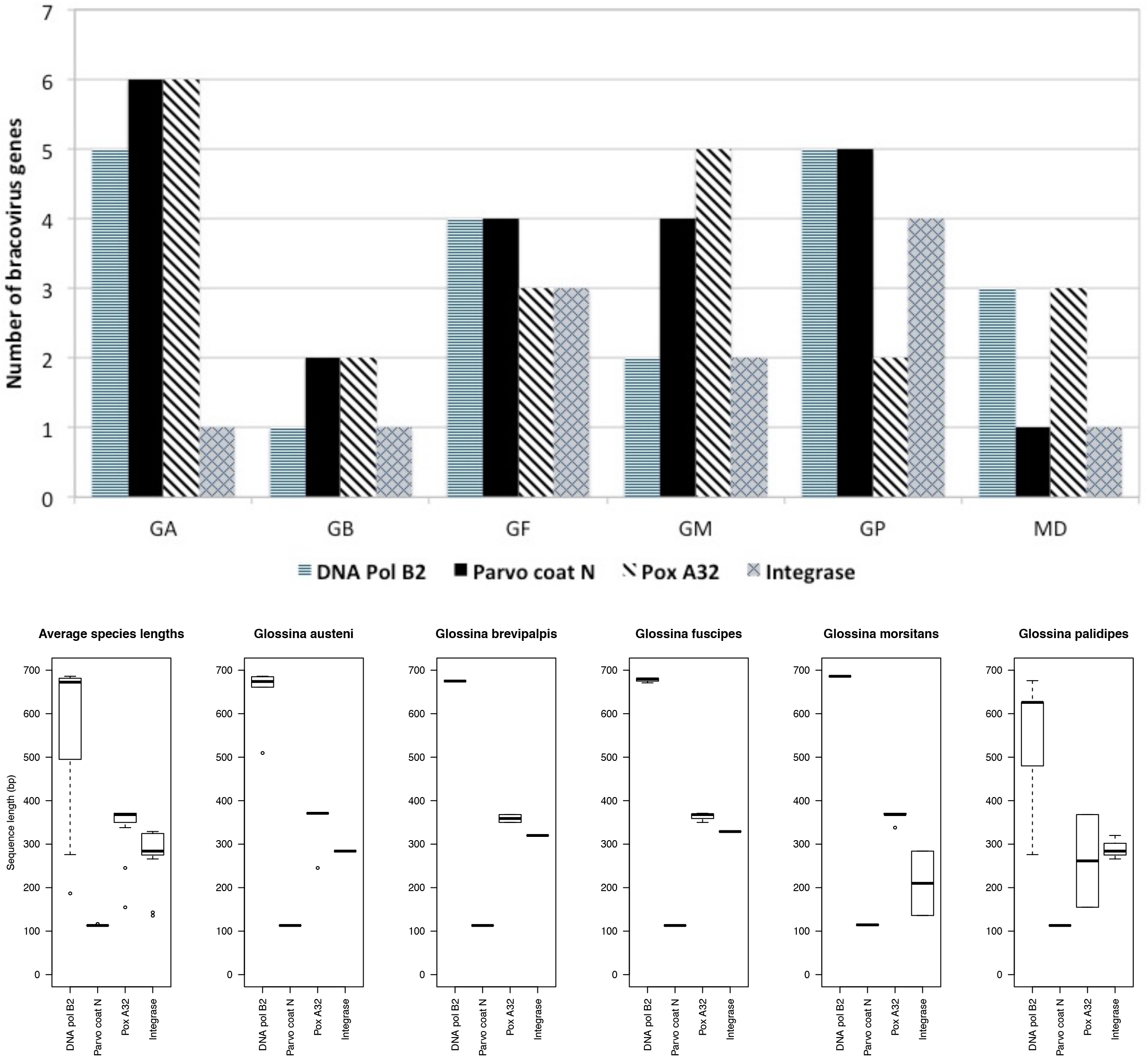
Differences in genomic bracovirus distribution and variation of sequence length in *Musca* domestica and *Glossina* species. (A) The distribution of Maverick associated genes identified by BLASTP searches (E values <1e-50) in *Glossina austeni* (GA), *G. brevipalpis* (GB), *G. fuscipes* (GF), *G. morsitans* (GM), *G. pallidipes* (GP), and *Musca domestica* (MD) genomes. *G. austeni* has the most number of Maverick associated genes (n=29) and *G. brevipalpis* the least (n=6). (B) Box plots depicting the entire range of gene lengths and the average gene length for each of the Maverick associated gene: DNA polymerase B2, parvo coat N, pox A32 and integrase. The plots show the average length for each gene across all the species, and within each species. The parvo coat N gene has the most conserved length across all species

### Phylogeny of Dipteran bracoviral sequences identifies two major clades

Phylogenetic reconstruction of each of the four Maverick associated *Dipteran* gene loci alongside orthologs from parasitoid wasps (*Cotesia congregata, Glyptapanteles flavicoxis* and *Nisonia vitripennis*) and a beetle (*Tribolium castaneum*) identified two major bracoviral clades, designated clade 1 and 2 respectively (Figure 2A). Interestingly, Dipteran sequences within each of the two clades consistently cluster together across all four loci implying similarity in direction of selection pressure. There are notable differences in the branching times of the clades for each gene locus. Clade 1 is more recent for DNA polymerase B2 and parvo coat N compared to the retroviral-like integrase and the poxvirus A32 protein. The opposite is true for clade 2. We evaluated the direction and magnitude of natural selection acting on these loci using the dN/dS ratio. We observe that the genes are under varying magnitudes of purifying selection (Table 1). Natural selection at the flanking sequences suggesting that Maverick associated genes were inserted at conserved regions of the host’s genome (Supplementary Figure 1). We did not observe terminal inverted repeats, which are associated with transposable elements.

**Figure 2.**
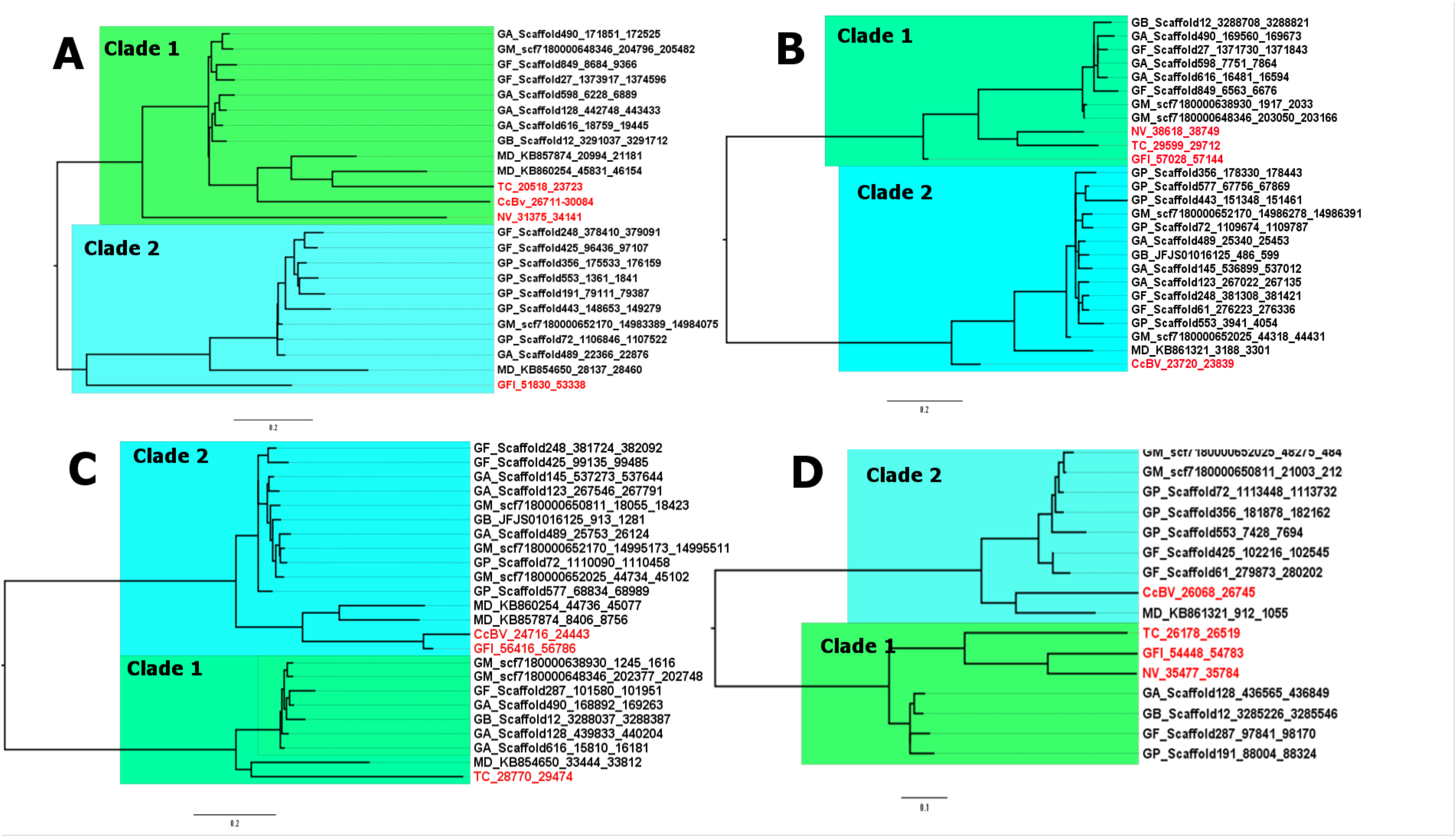
Phylogenetic reconstruction Maverick associated genes. Trees for each gene locus were reconstructed as follows: (A) DNA polymerase B2 genes, (B) parvo coat N genes, (C) pox A32 and (D) integrase. Orthologs at each loci separated in to two distinct clades (clade 1 and 2). Sequences that clustered together in each clade are highlighted in blue and green. Orthologs from parasitoid wasps (*Cortesia congregate* (CcBV), *Glyptapanteles flavicoxis* (GFl) and *Nisonia vitripennis* (NV)) and a beetle (*Tribolium castaneum* (TC)) are highlighted in red.

**Table 1.** The direction and magnitude of selection pressure on Maverick associated genes. Shows the magnitude and direction per clade for a given loci, and the overall loci dN/dS values for all the analysed bracoviral sequences.

| Gene | Clade 1 | Clade 2 | Overall |
| --- | --- | --- | --- |
| DNA Pol B2 | 0.85 | 1 | 0.66 |
| Parvo coat N | 0.18 | 0.94 | 0.77 |
| Pox A32 | 0.78 | 0.16 | 1.1 |
| Integrase | 1.21 | 0.17 | 0.85 |

### Detection of Maverick sequences within recent African tsetse flies

We extracted genomic DNA from a collection of male and female tsetse flies from Kenya (*G. pallidipes, G. brevipalpis* and *G. fuscipes*) and Malawi (*G. m. morsitans*). We amplified Maverick genes in all tsetse fly species. Our results confirm that the annotated Maverick genes exist in tsetse flies collected from different geographic locations (Figure 3). The products obtained were of the expected sizes (Supplementary Table 2).

**Figure 3.**
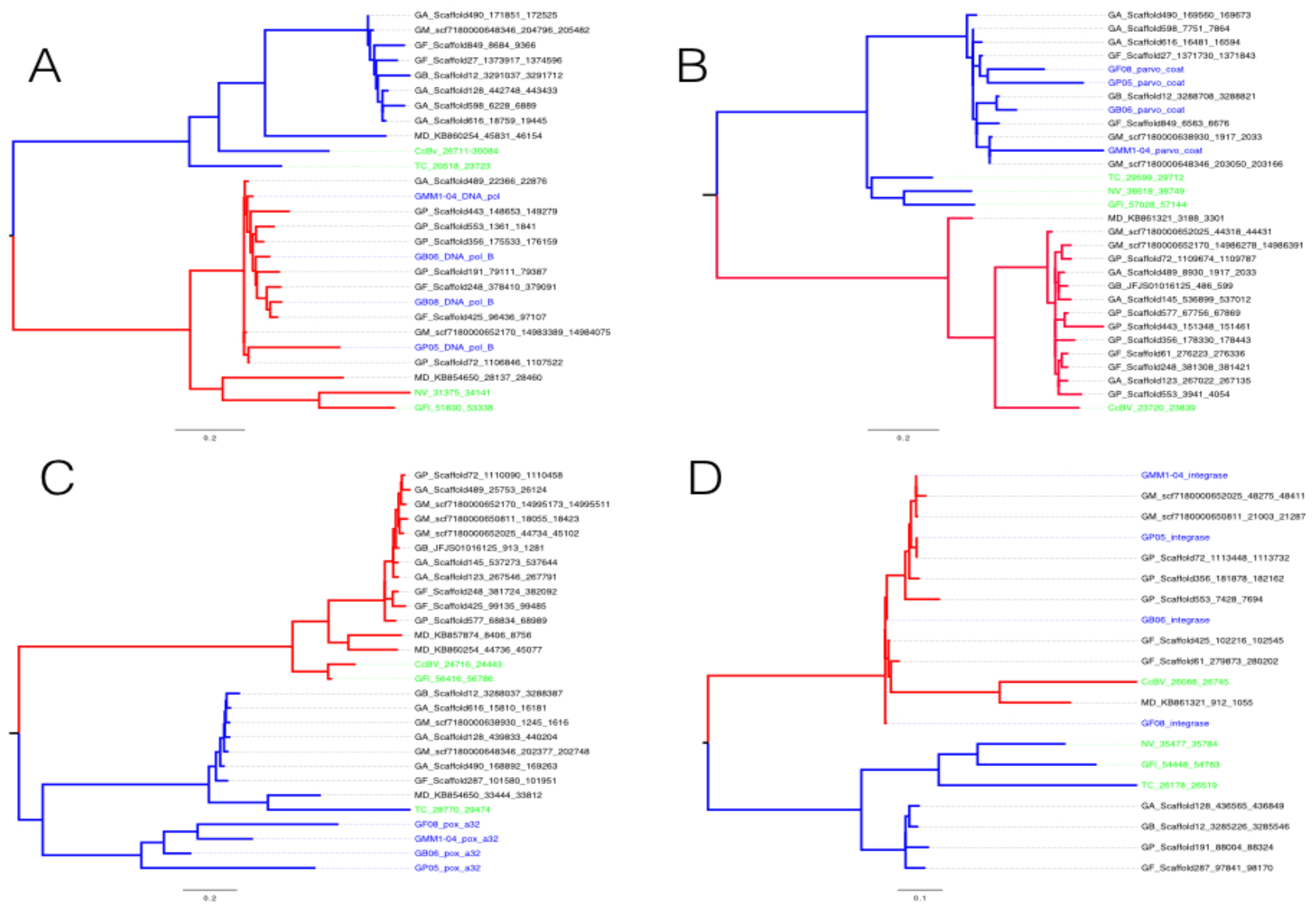
Detection of Maverick sequences within recent African tsetse flies. Trees for each of the four genes: (A) DNA polymerase B2 genes, (B) parvo coat N genes, (C) pox A32 and (D) integrase identified in the Glossina genomes (black) were reconstructed together with sequences generated from tsetse flies recently collected in Kenya and Malawi (blue) and reference sequences (green).

## Discussion

Endogenous bracoviral sequences have been identified in the genomes of parasitoid wasps and *Glossina morsitans morsitans* (International Glossina Genome Initiative 2014). Bracovirus present a unique symbiotic relationship between eukaryotes and endogenous viruses. Understanding their organisation significantly contributes new vector control strategies and the development of vectors for gene therapy (Bezier et al. 2009). For example, the polydnavirus *Oryctes rhinoceros nudivirus* (OrNV) has been used as a biological control agent in palm tree farming against rhinocerous beetle (Bezier et al. 2009). We show the prevalence of polydnaviruses (PDVs) in *Glossina*, provide evidence that they are descended from a single ancestor after initial host integration, and their presence in the reference *G. m. morsitans* is not a single random genetic introgression event. We also show that PDVs vary in size, but it is still unclear whether they undergo intra-genomic rearrangements, in their host genomes.

We identified four PDV genes of Maverick origin: DNA polymerase B2 involved in DNA excision repair and initiation of replication (DNA Pol B 2), the N terminal region of the parvovirus coat protein VP1 (Parvo coat N) that is important for virion retention and transduction by insect vectors, a retroviral-like integrase, and poxvirus A32 protein (Pox A32) that encodes an ATPase involved in virion DNA packaging. Although bracoviruses in wasps are co-opted to ensure their successful reproduction, their role in *Diptera* that do not share this mode of reproduction remains unclear. We speculate that they are redundant in tsetse flies. We did not identify a protease gene confirming that the *Dipteran* Maverick has lost the viral maturation ability but retains the transposition ability due to the presence of the integrase (Pritham et al. 2007). Maverick-associated protease genes, which play key roles in successful reproduction, are present in the bracoviral genomes of parasitoid wasps (Dupuy et al. 2011). *G. austeni* has the most abundant number of Maverick genes, and *G. brevipalpis* the least number, and the palpalis group is intermediate. Phylogeny reconstruction of the *Glossina* species has shown that fusca group (*G. brevipalpis*) forms the deepest branches, followed by morsitans (*G. pallidipes, G. austeni* and *G. morsitans*) and palpalis (*G. fuscipes*) groups (Chen et al. 1999). This implies that *G. brevipalpis* is the least diverse species and has fewer genome rearrangement events.

Homologous PDV sequences identified across the 5 *Glossina* species and *M. domestica* provide evidence that a single integration event of the viral genome occurred before the evolutionary radiation of different insect orders. Subsequent PDV genetic diversification occurs as they co-evolve with the host insect genome. Our findings improve knowledge on PDVs, and explain presence and retention in tsetse flies, yet tsetse flies do not exhibit parasitoid reproduction. It is unclear when exactly parasitoid wasp and *Dipteran* PDVs separated and diversified in their different hosts. Our findings support previous findings that bracoviruses are descended from a common ancestor in the Paleozoic Era, and raise the possibility of integration of PDVs before the separation of *Hymenoptera, Coleoptera, Lepidoptera* and *Diptera* (Theze et al. 2011).

Mavericks (also known as polintons) are a group of transposable elements, present in most eukaryotes - including parasitoid wasps (Dupuy et al. 2011), which combine features of *bona fide* viruses and transposable elements (Koonin et al. 2015). They can subsist as viruses or transposons depending on the combination of genes present (Koonin et al. 2015). Mavericks are thought to be the first group of double-stranded DNA viruses to evolve from bacteriophages (Krupovic and Koonin 2015). They continuously evolve during vertical eukaryotic host transmission, and are actively involved in shaping the genomes and biology of eukaryotes by acting as ‘epicenters’ of gene accretion and genome expansion (Krupovic and Koonin 2015).

We observed branching of the Maverick genes into two distinct clades with notable differences in the branching times, for all four groups of Maverick genes. This is consistent with previous findings that used different phylogenetic and taxonomic sampling methods to highlight two types of Maverick sequences. The first type is thought to replicate in the mitochondrion, and second in the cytoplasm (Krupovic and Koonin 2015). We show the co-occurrence of two types of Maverick genes in each *Dipteran* genome. We speculate that lineage specific duplications may have resulted in diverse sets of genes under different magnitudes of evolutionary pressures giving rise to the two clades. Previous findings have shown that multiple copies of homologous PDVs have been identified in wasps at different loci of the insect genome (Desjardins et al. 2008; Bezier et al. 2013). This evolution of endogenous viral genes involves numerous rearrangements resulting from successive lineage-specific duplications, each creating genetic variants (Bezier et al. 2013).

Terminal inverted repeats (TIRs) were absent in the flanking regions of the Dipteran Mavericks. TIRs have also been shown to be absent in some parasitoid wasp bracoviral genomes, for example the *Cortesia congeregata* bracoviral genome (CcBV) (Dupuy et al. 2011). This absence has been ascribed to cumulative insertion and deletion events that blur genetic identification, and these events have minimal effects in the host’s fitness (Langley and Charlesworth 1997). Transposable elements without TIRS are dysfunctional and are degraded. However, the retention of *Dipteran* Maverick genes suggests that they are under purifying selection, albeit with unknown function (Desjardins et al. 2008). This group of double-stranded DNA has however still accumulated variation over time making it challenging to reconstruct macro-evolutionary history of viruses beyond 100 Mya (Theze et al. 2011).

We examined the direction and magnitude of selection on the Maverick genes. Overall; these genes are under purifying selection, except pox A32, which is under positive selection. This is surprising because PDVs in *Diptera* appear to be non-functional viral fragments that are evolutionary dead ends. We also examined selection pressure of host genome sequences flanking these loci to determine whether the bracoviral genes were inserted in conserved regions. The flanking regions were relatively conserved; with the exception of the sequence upstream of parvo coat N, and downstream of the integrase genes. Similarity in the magnitude and direction of selection pressure suggests that bracoviral genes have lowered functional constraints to globally co-evolve with the host genome (Herniou et al. 2013).

The presence of bracoviruses in tsetse flies collected in different parts of Africa (Kenya and Malawi) compared to the reference tsetse fly genomes indicates that the genes are widely distributed and stably associated with *Diptera*. This study provides evidence that this was neither a result of a pathogenic virus that contaminated the reference tsetse fly genome, nor a single case of being parasitized by bracoviruses of wasp origin. We also assess the Maverick genetic diversity, from widely geographically distributed insect hosts.

### Conclusion

We conclude that bracoviruses are present in *Diptera* and share close homology. These bracoviral sequences can be grouped into two major groups, are under purifying selection, and co-evolve with the host genome. Approaches that could exploit bracoviruses for vector control remain to be determined.

## Materials and Methods

### Identification of bracoviral domain sequences

Bracoviral related sequences (n=310) present in the *G. m. morsitans* proteome, previously described by the International Glossina Genome Initiative (International Glossina Genome Initiative 2014), were retrieved using a Perl script. These sequences were subjected to BLASTP and DELTA BLAST searches against NCBI’s Conserved Domain Database (CDD) (28 May 2015 release) (Marchler-Bauer et al. 2011) to identify bracoviral related domains [4]. Identified domains were also confirmed by querying the sequences against Pfam-A.hmm (13 March 2013 release) downloaded from the Pfam 27.0 (Finn et al. 2014) protein families database. Searches against the Pfam-A.hmm were accomplished using the hmmscan command (cut off e-value of <1.0e-10) in HMMER version 3.1b1 (Eddy 2011).

The identified bracoviral domain sequences were retrieved and used as query to find homologs in the *G. austeni, G. brevipalpis, G. f. fuscipes, G. m. morsitans, G. pallidipes* and *M. domestica* protein datasets retrieved from VectorBase (Lawson et al. 2009) using the tBLASTn algorithm (cutoff e-value <1.0e-10 and BLOSSUM62 scoring matrix) (Giraldo-Calderon et al. 2015). The identified bracoviral domain sequences were verified by alignments against NCBI’s non-redundant protein database using the BLASTX algorithm. These domain sequences were extracted and used for further analysis.

### Phylogenetic reconstruction of bracoviral domains

DNA Pol B2, parvo coat N, pox A32, and an integrase-like domain sequences from *Glossina spp*. (*G. austeni, G. brevipalpis, G. f. fuscipes, G. m. morsitans* and *G. pallidipes*), *Musca domestica*, and orthologs from parasitoid wasps (*Cotesia congregata, Glyptapanteles flavicoxis* and *Nasonia vitripennis*) and a beetle (*Tribolium castaneum*) (Dupuy et al. 2011), were used for phylogeny reconstruction. Domains were aligned using Multiple Sequence Comparison by Log Expectation (MUSCLE) (Edgar 2004). Maximum likelihood (ML) phylogenetic analysis of the multiple aligned sequences with bootstrap values of 100 replicates was performed using PHYML version 3.5 (Guindon and Gascuel 2003).

### Magnitude and direction of selection pressure

The magnitude and direction of selection pressure on the bracoviral domain sequences was tested based on the ratio (ω = d_N_/d_S_) of the average number of non-synonymous substitutions per non-synonymous site to the average number of synonymous substitutions per synonymous site (d_S_). If ω = 1, amino acid substitution is assumed to be under neutral selection, ω > 1 is indicative of positive selection whereas ω < 1 is evidence of negative or purifying selection. A sequence alignment was generated for each of the loci from the *Glossina* species and *Musca* domestica domain sequences; as well as sequences from parasitoid wasps (*Cotesia congregata, Glyptapanteles flavicoxis* and *Nasonia vitripennis*) and a beetle (Tribolium castaneum. Each alignment was then uploaded to the SNAP program (Korder 2000) (www.hiv.lanl.gov) that calculates synonymous and non-synonymous substitution rates to determine the magnitude of selection pressure. Sequences (length = 50bp) flanking either side of each domain sequence were also extracted using Perl scripts and the magnitude of selection pressure examined. This was similarly evaluated for each clade per loci under examination.

### Laboratory validation

Laboratory validation of the four gene loci was performed using conventional polymerase chain reaction (PCR) amplification. A list of primers used is provided in Supplementary Table 2. Insect DNA was extracted using DNeasy Blood & Tissue Kit (Qiagen, Hilden, Germany), using the manufacturer’s instructions. Briefly, initial DNA template denaturation for 3 minutes at 95^0^C, followed by 40 cycles of 30 seconds denaturation at 95^0^C, 30 seconds of annealing (55^0^C for DNA Pol B2, 51^0^C for Parvo coat N, 54^0^C for Pox A32 and 55^0^C for integrase), and 30 seconds of elongation at 72^0^C. The final extension was 7 minutes at 72^0^C. The reaction mix contained 5 µl Dreamtaq mastermix 2x, 1 µl (10 pmol) of each primer, topped up to 10µl with PCR water. Sanger sequencing on both strands of the PCR products was outsourced from Macrogen (Seoul, South Korea). The sequences obtained were processed and compared with the original sequences to determine genetic diversity.

### Data access

The gene sequence data generated in this study is publicly available on GenBank using the assigned accession numbers KU925563-KU925578.

## Acknowledgments

This work was supported by Training Health Researchers into Vocational Excellence in East Africa (THRiVE), grant number 087540 funded by the Wellcome Trust. Its contents are solely the responsibility of the authors and do not necessarily represent the official views of the supporting offices. BWK received a pump-priming grant from THRiVE.

## Author contributions

BWK and DM conceived and designed the study. KMK, MFA, WO, BWK performed experiments and analysis. EM provided samples. BWK and KMK wrote the initial manuscript draft. KMK, MAF, WO, EM, DM and BWK wrote the final version of the manuscript.

## Disclosure declaration

The authors declare no conflicts of interest.

## Supplementary Figure Legends

**Supplementary Table 1. The distribution of Maverick associated genes in different Dipteran species genomes**. The description is as follows: species abbreviations, genome scaffold, and start and end base pair position for each gene. The abbreviations describe genes from the six different species: *Glossina austeni* (GA), *Glossina brevipalpis* (GB), *Glossina fuscipes* (GF), *Glossina morsitans* (GM), *Glossina pallidipes* (GP), and *Musca domestica* (MD). Genes present on the same scaffold are in the same row. The stars indicate the number of genes present per scaffold, and if all four Maverick associated gene loci are present are highlighted in red.

**Supplementary Table 2: List of Genes, Primer sequences and the expected product size. ‘**For’ stands for Forward primer and ‘Rev’ stands for reverse primer.

**Supplementary Figure 1. Magnitude and direction of selection pressure at Maverick associated loci and their flanking regions**

**Supplementary Table 2**

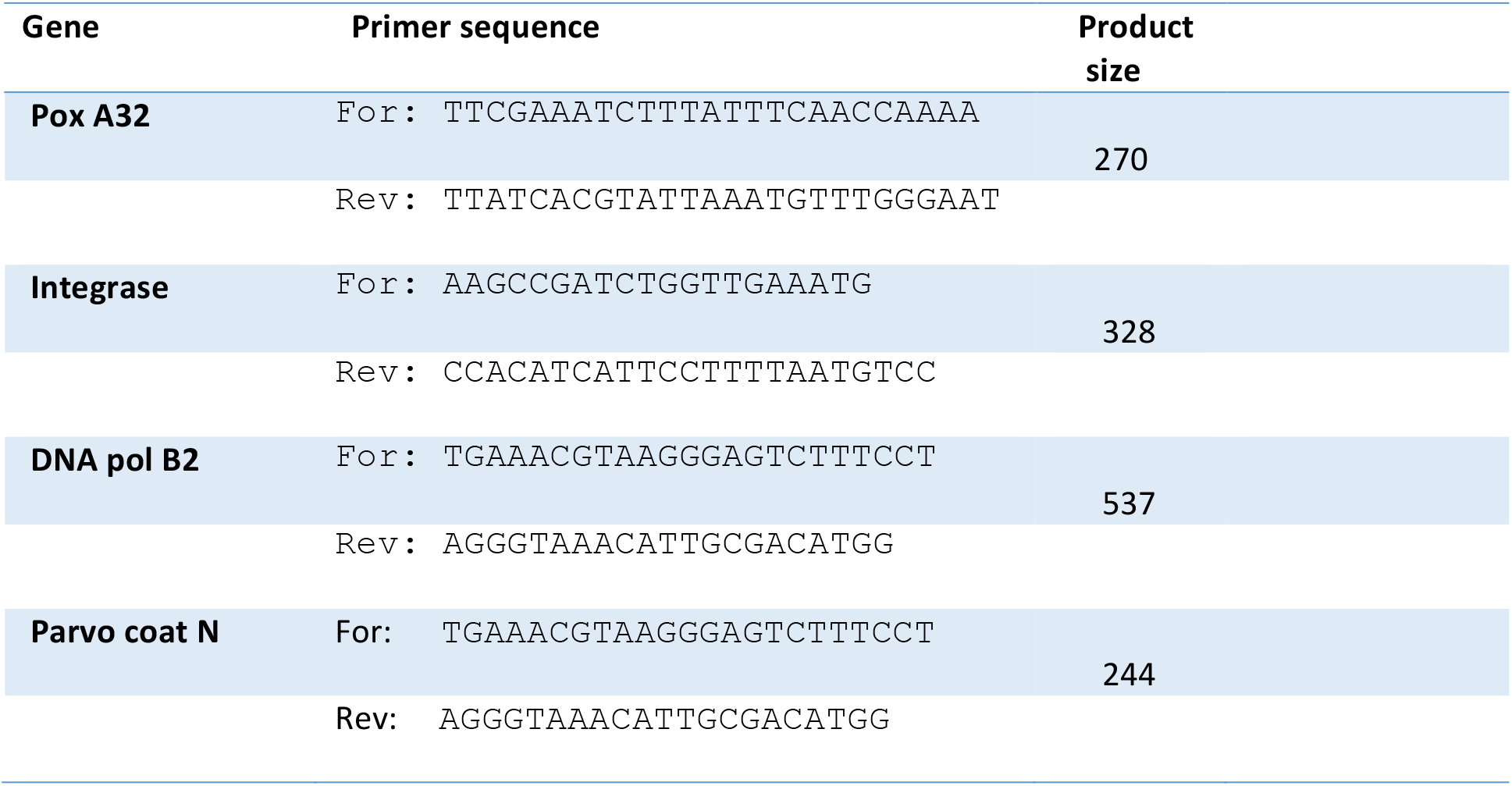

